# Levitational Cell Cytometry for Forensics

**DOI:** 10.1101/2020.11.09.374843

**Authors:** Deniz Yagmur Urey, Hsi-Min Chan, Naside Gozde Durmus

## Abstract

Here, a method for label-free, real-time interrogation, monitoring, detection and sorting of biological rare cells in magnetically-suspended heterogeneous samples is developed. To achieve this, heterogeneous populations of cells were levitated and confined in a microcapillary channel. This strategy enables spatiotemporal differential magnetic levitation of rare fragile dead cells equilibrating at different heights based on the balance between magnetic and corrected gravitational forces. In addition, sorting of fragile rare dead cell populations is monitored in real-time. This technique provides a broadly applicable label-free tool for high resolution, real-time research, as well as forensic evidence processing of rape kits. This method is validated with forensic mock samples dating back to 2003, isolating sperm from epithelial fraction with >90% efficiency and >97% purity. Overall, this method reduces the processing time by over 20-fold down to 20 minutes, eliminating centrifugation and labels, and providing an inexpensive and a high-yield alternative to the current centrifuge-based differential extraction techniques. It can potentially facilitate the forensic downstream genomic analyses, accelerating the identification of suspects, and advancing public safety.

## 1. Introduction

35% of women worldwide have been a victim of sexual assault in their lifetime. However, as a result of the time-consuming, centrifugation-based and laborious differentiation of the perpetrator’s cells from the victim’s cells, processing the evidence to identify the perpetrator has been limited. Moreover, 60-90% of the perpetrator’s sperm in forensic kits is lost during the process. Analyzing evidence after cases of sexual assault in a timely manner is crucial for the identification of the perpetrator. Yet, the drawbacks in the protocols create a major problem for the criminal justice system, and more importantly, for the victim. According to Rape, Abuse & Incest National Network, an investigative report in 2015 identified 70,000 sexual assault kits from over 1,000 police departments that were not tested for DNA evidence. This backlog is due to the lack of sufficient resources and it takes a long time to obtain results to test, process, and profile samples in crime labs^[1]^.

For the examination of forensic kits, the epithelial cells of the victim first have to be separated from sperm of the perpetrator before DNA analysis. One of the major drawbacks has been the requirement of high concentration of sperm from the swab sample because up to 90% of sperm is lost through the current methodology during examination. This is caused by the time-consuming (up to 8 hours), labor-intensive steps of selective cell lysis, centrifugation, and separation into female and male cell fractions for the methods used in forensics.^[2]^ There have been several attempts to establish alternative methods for the differential extraction of sperm, such as acoustic trapping^[3]^, laser microdissection^[4]^, magnetic bead-based separation^[5, 6]^, microfluidic capture based separation^[7, 8]^, and antibody-based sperm capture^[6]^. However, these technologies have not been utilized in practical applications, due to factors such as sperm detection/sorting efficiency, assay time, and usage of label markers (Table S1). In particular, the antibody-based sperm extraction methods have difficulties to work with aged samples due to the changes in the antigen specificity of sperm over time, making antibodies less capable to bind to sperm, and thus limiting their utility for forensic samples^[5, 7]^.

Here, in order to address the sperm separation efficiency and test limitations around assay duration, we have utilized the inherent differences in cellular content of sperm and epithelial cells through the principles of magnetic levitation^[9]^ (**Figure 1**). We have developed a levitation-based method for differential extraction of cells in a collected biological sample and spatiotemporal separation of sperm and epithelial cells by leveraging differences in levitation due to density and paramagnetic medium uptake. This approach enables to precisely sort rare sperm in concentrations as low as 100 sperm per sample, working with extremely small sample volumes as low as 30 μL (typical in forensic samples); to reduce the differential extraction time from 8 hours to 20 minutes; to achieve an unprecedented high sperm detection and sorting efficiency, which is not possible with the current methods. The workflow of our levitation setup can be seen in Figure 1a and the sorting capability is introduced uniquely to handle small and rare samples in Figure 1b.

**Figure 1.**
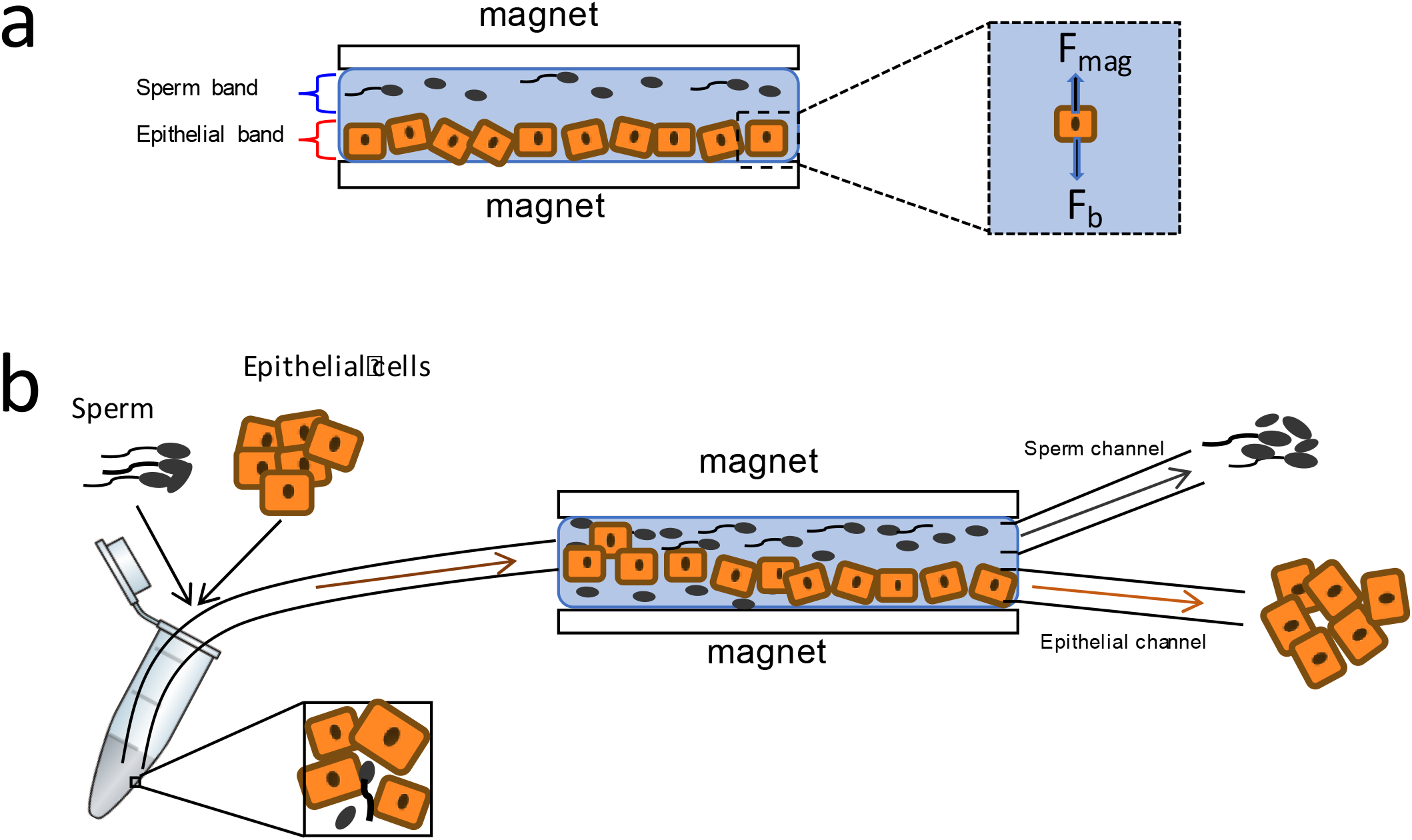
Workflow of differential extraction of sperm from epithelial cells using magnetic levitation. **a)** Sperm are spiked in epithelial cells, and the mixture is loaded to the capillary channel of the magnetic levitation device. Magnetic fields and gradients counteract gravity. Both cell types have a different levitation profile, based on their densities. Sperm levitate higher than epithelial cells, and the cells form levitation bands within the channel. **b)** Spiked sperm and epithelial mixture is loaded to the capillary channel. Once sperm and epithelial cells form their levitation bands, sperm can be collected from the top channel (sperm channel), and epithelial cells can be collected from the bottom channel (epithelial channel).

## 2. Results

### 2.1. Magnetic Levitation of Sperm and Epithelial Cells

In levitation, cells are diamagnetic^[10]^ and they are suspended in a biocompatible, paramagnetic medium using forces generated by magnetic fields and gradients counteracting gravity^[9]^. We present that the separation of sperm from epithelial cells is possible due to differences in the levitation heights of these cells in forensic samples. As all cells have different densities and all organic materials are diamagnetic, magnetic levitation and density gradient based separation of single cells is used to separate sperm from epithelial cells. The magnetic susceptibility difference between a cell and its surrounding paramagnetic medium causes it to move away from a higher (i.e., close vicinity at the magnets) to a lower magnetic field strength site (i.e., away from the magnets) until gravitational, buoyancy and magnetic forces acting on the cells reach an equilibrium. Cells are levitated at a final position between the two magnets, where the magnetic force (Fmag) equals the buoyancy force (Fb). As the cells flow, the platform monitors the equilibrium heights of each individual cell in real-time, magnetically focuses and separates sperm and epithelial cells at different levitation bands, based on their unique magnetic signatures (More information on the underlying mechanisms of levitation of cells in the microcapillary is provided in Supplementary Information). In order to quantify the average levitation heights of the two cell types, we first performed experiments levitating sperm and then epithelial cells separately in channels (Figure 2a). We utilized the magnetic levitation device consisting of two permanent magnets with the same poles facing each other. The tilted side mirrors were used to simply integrate the tool to any microscope and measure the levitation heights of cells inside a capillary channel. The use of a channel enabled to easily load and unload the samples given the necessity to work with extremely small volumes in forensic samples. For instance, these sample can be extracted from a single cotton swab emphasizing the need for the device to work with small sample volumes 10-50 μL and very small number of sperm mixed in a large epithelial cell (E-cell) population.

**Figure 2.**
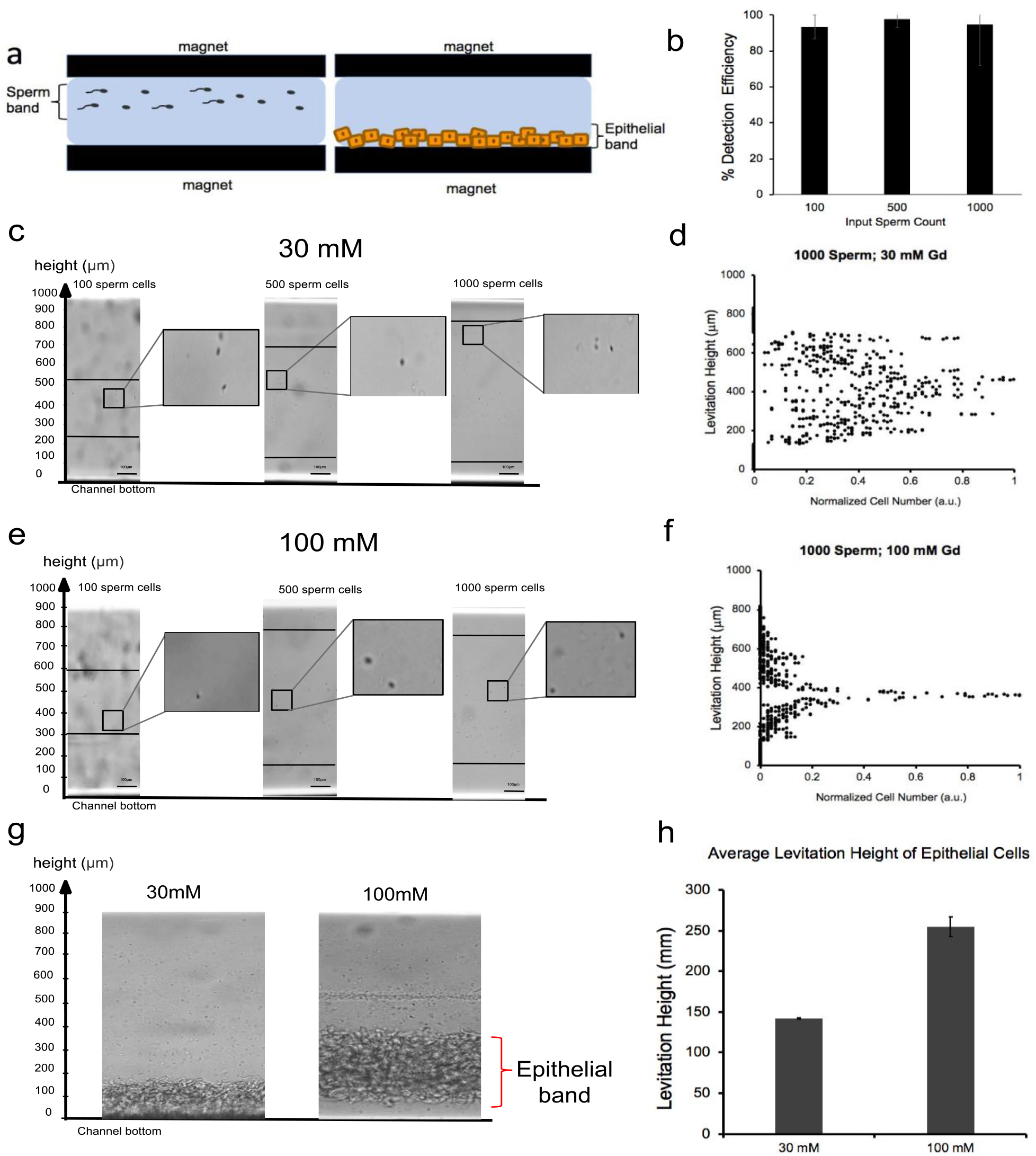
Magnetic levitation of sperm and epithelial cells in separate devices. **a) a)** Schematic representation for comparing the levitation profiles of sperm and epithelial cells. 100, 500, and 1000 sperm cells/sample volume (30 μL) was loaded in the capillary channel with 30 mM and 100 mM paramagnetic medium. Sperm cells reached their levitation heights after 15-20 minutes. On the other hand, 10,000 and 50,000 epithelial cells were loaded in the capillary channel with 30 mM and 100 mM paramagnetic medium. Epithelial cells reached their levitation height after 4-7 minutes. **b)** Sperm detection efficiency was calculated for 100, 500, and 1000 sperm in the capillary channel. Sperm detection efficiency is defined as the ratio of the number of sperm detected in the capillary channel and the number of sperm initially loaded to the channel. **c)** 100, 500, and 1000 sperm/sample is levitated in 30 mM paramagnetic medium. The lines indicate the levitation bands. Width of the levitation band expands as sperm number increases. **d)** Levitation height vs. normalized cell number graph for 1,000 sperm with 30 mM paramagnetic medium is plotted according to each sperm’s levitation profile. **e)** 100, 500, and 1,000 sperm cells/sample was levitated in 100 mM paramagnetic medium. The horizontal lines mark the levitation bands. Width of the levitation band expands as the sperm number increases due to increasing heterogeneity. The levitation height was higher than that of in 30 mM paramagnetic medium. **f)** Levitation height *vs*. normalized cell number graph for 1,000 sperm in 100 mM paramagnetic medium is plotted. A clearer peak in average levitation height is observed at 100 mM paramagnetic medium. **g)** 10,000 and 50,000 epithelial cells are levitated with 30 mM and 100 mM paramagnetic medium. **h)** Average levitation height of 10,000 epithelial cells is plotted for 30 mM and 100 mM paramagnetic medium, according to cell levitation profiles. The epithelial cells have a lower levitation height in 30 mM paramagnetic medium than that in 100 mM paramagnetic medium.

First, to test the limitations of our method, we used very low sperm counts down to 100 sperm. In sperm levitation experiments, we used two main parameters: i) sperm counts of 100, 500, and 1,000; ii) non-ionic paramagnetic medium (gadolinium) concentrations of 30 mM (Figure 2c, d) and 100 mM (Figure 2e, f). A total sample volume of 30 μL was used for each trial. We showed that the method yields high detection efficiencies for 100, 500, and 1000 sperm/sample volume as 93.3%, 97.9%, and 92.8%, respectively (Figure 2b, Table S2, Supporting Information). The 100 mM paramagnetic medium levitated the sperm into a tighter levitation band focused by the stronger magnetic field forces (Figure 2f, Video S1, Supporting Information).

Second, we asked whether epithelial cells can be levitated, and we performed experiments with two main parameters: i) epithelial cell counts of 10,000 and 50,000; ii) paramagnetic medium concentrations of 30 mM and 100 mM (Figure 2g). The levitation process took 7 minutes for 100 mM; and 5 minutes for the 30 mM paramagnetic medium to come to a final equilibrium in levitation due to stronger magnetic forces in the 100 mM medium case against the gravity. We observed that the levitation height of epithelial cells in 30 mM paramagnetic medium was lower than that in 100 mM concentration, which later enabled us to optimize the epithelial cell levitation relative to the sperm levitation for the maximum spatial separation (Figure 2g, h).

### 2.2. Magnetic Levitation of Sperm and Epithelial Cell Mixtures

After identifying the relative levitation height ranges for individual cell types, we investigated the levitation profiles in heterogeneous mixtures of epithelial cells and sperm that represent forensic samples from rape kits (Figure 3a). Yet, the concentration difference of epithelial cells and sperm is very high within the sample, due to the fact that the swab is taken from the body of the victim. To mimic a rape forensic kit sample, we spiked a low number of sperm cell into a high number epithelial cell sample. We observed that the epithelial cells in mixture obtained their levitation bands by sinking to the bottom within 4-7 minutes in 30 mM paramagnetic medium (Figure 3b, Video S2, Supporting Information). This process leaves the rest of the top channel for sperm to levitate and come to an equilibrium. Since one of the current drawbacks of the existing methods is the loss of sperm during differential extraction^[7]^, we gradually reduced the number of sperm spiked into epithelial cells between each set of experiments to be able to see whether a high separation efficiency could be maintained even with very low sperm counts. We tested three main parameters: i) sperm cell count of 100, 500, and 1000 per sample; ii) epithelial cell count of 10,000 and 50,000 per sample; iii) paramagnetic medium concentrations of 30 mM and 100 mM. We optimized the epithelial cell count to be 10,000 cells per 30 μL of sample, where the separation accuracy is the highest at this concentration (Figure 3c). The lower number of epithelial cells in the channel also allowed a thinner band for epithelial cells reducing the likelihood that some sperm may be stuck in the thicker E-cell bands as cells levitate in the channel (Figure S1, Supporting Information). This process reduces the overall assay time required for levitation. The separation efficiencies for 100, 500, and 1000 sperm were 93.9%, 96.1%, and 94.5% respectively (Figure 3d, Table S3, Supporting Information). At 30 mM, we observed that sperm had a wider, more dispersed levitation band compared to that at 100 mM. On the other hand, levitation at 30 mM concentration yielded a higher separation efficiency, as the epithelial cells had a lower levitation height that increases the space in the channel dedicated to sperm levitating above the epithelial cell band. This also facilitates the later sorting step as the sperm do not need to reach an equilibrium for them to be collected given that the larger sized epithelial cells reach the bottom within 4-7 minutes. We have shown that the epithelial cells equilibrate at a higher levitation height when 100 mM paramagnetic medium concentration is used. Thus, we can conclude that the levitation height of epithelial cells at the bottom of the channel under 30 mM paramagnetic medium concentration is not solely due to the gravity but is rather caused by magnetic forces in the levitation device.

**Figure 3.**
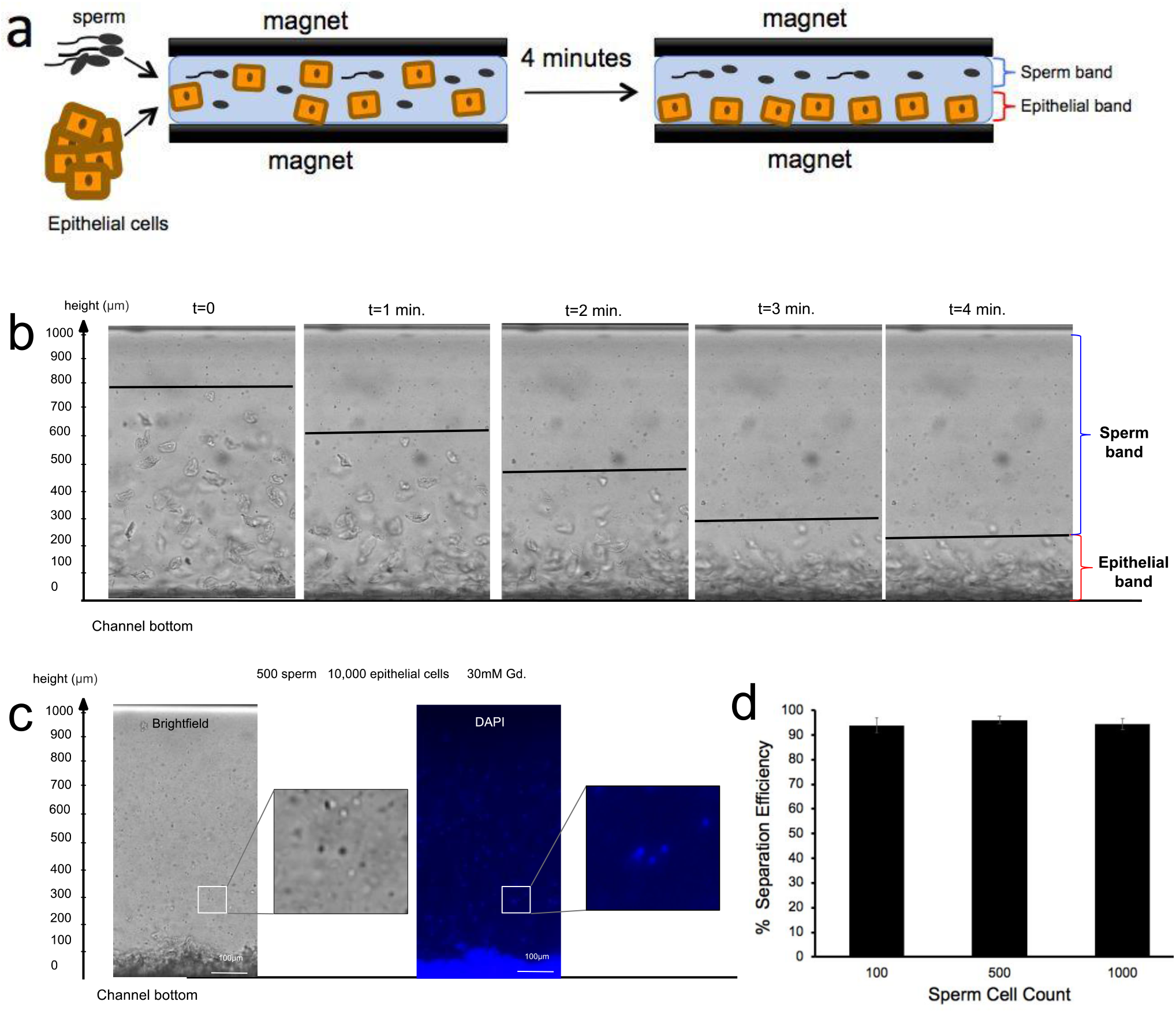
Magnetic levitation and separation of sperm from epithelial cells. **a)** Schematic representation of the experimental workflow. 100, 500, and 1,000 sperm/sample volume were spiked in 10,000 epithelial cells in 30 mM paramagnetic medium. 30 μL total sample volume was loaded into the capillary channel. Sperm levitated at a higher levitation band, whereas the epithelial cells sank down to the bottom of the capillary channel. **b)** Real-time monitoring and imaging of separation of sperm from the epithelial cells. Epithelial cells settled to the bottom of the capillary channel 4 minutes after the sample was loaded into the capillary. An image was taken after each minute to show their change in levitation height over time. Gradual decrease in levitation height of epithelial cells is shown with lines. Epithelial cell band and sperm band are shown in the final image. **c)** The levitation profiles of sperm and epithelial cells are imaged under bright field and DAPI. Sperm and epithelial cells can be detected as fluorescent blue because of DAPI staining. **d)** The separation efficiency of sperm cells in 10,000 epithelial cell population was calculated for 100, 500, and 1000 sperm cells/sample volume. Separation efficiency was defined as the ratio of the number of sperm counted above epithelial cell band and the total number of sperm in the channel. The sperm can be separated from the epithelial fraction with high efficiency; such as 93.9%, 96.1%, and 94.5% for 100, 500, and 1,000 sperm, respectively.

### 2.3. Sorting of Sperm from Epithelial Cells in Magnetic Levitation Platform

After the spatiotemporal separation of sperm from epithelial cells in the magnetic levitation system was optimized, we investigated methods to recover the spatially separated cells by a microfluidic sorting mechanism (Figure 4a). We performed flow-based sorting experiments in order to sort and isolate sperm from the top of a microfluidic capillary (Sperm channel), and epithelial cells from the bottom of the microfluidic capillary (Epithelial channel) (Videos S3-S4, Supporting Information). For the sorting experiments, the total sample volume was kept constant at 30 μL. As it is shown in earlier studies, the sensitivity and precision of levitation measurements increases at lower paramagnetic conditions [7]. Therefore, we decided to work with lower paramagnetic concentrations to process forensics samples in this study. Furthermore, since the epithelial band was farther from the sperm cells under 30 mM paramagnetic medium than it was under 100 mM paramagnetic medium, a better and clearer separation between the 2 cell types was detected under 30 mM paramagnetic medium (Figure 2g). Thus, a higher sorting accuracy was achieved. We experimented with 10,000 epithelial cells, 30 mM paramagnetic medium concentration, and i) 100, 500, 1,000 spiked sperm/sample volume. The epithelial cell counts and the paramagnetic medium concentration were kept constant, as they were optimized for the highest separation efficiency as described above. Here, we pre-stained the sperm with DAPI to be able to distinguish them from epithelial cells after mixing. After running the sample, we differentiated the sperm based on staining for the counting process. We imaged the spiked sample both before sorting (Figure 4b) and after sorting to evaluate the distribution of sperm and epithelial cells in the top (Sperm channel) (Figure 4c) and bottom collection channels (Epithelial channel) (Figure 4d). After 3 different trials with 10,000 epithelial cells and 100, 500, and 1000 sperm, we obtained an average sorting efficiency of 92.3%, 92.2%, and 93.6%, respectively (Figure 4e, Table S4). The sorting efficiencies were similar for all sperm counts, yet 100 sperm had the largest standard deviation, due to the low count of spiked cells in a limited sample volume (i.e., 30 µl).

**Figure 4.**
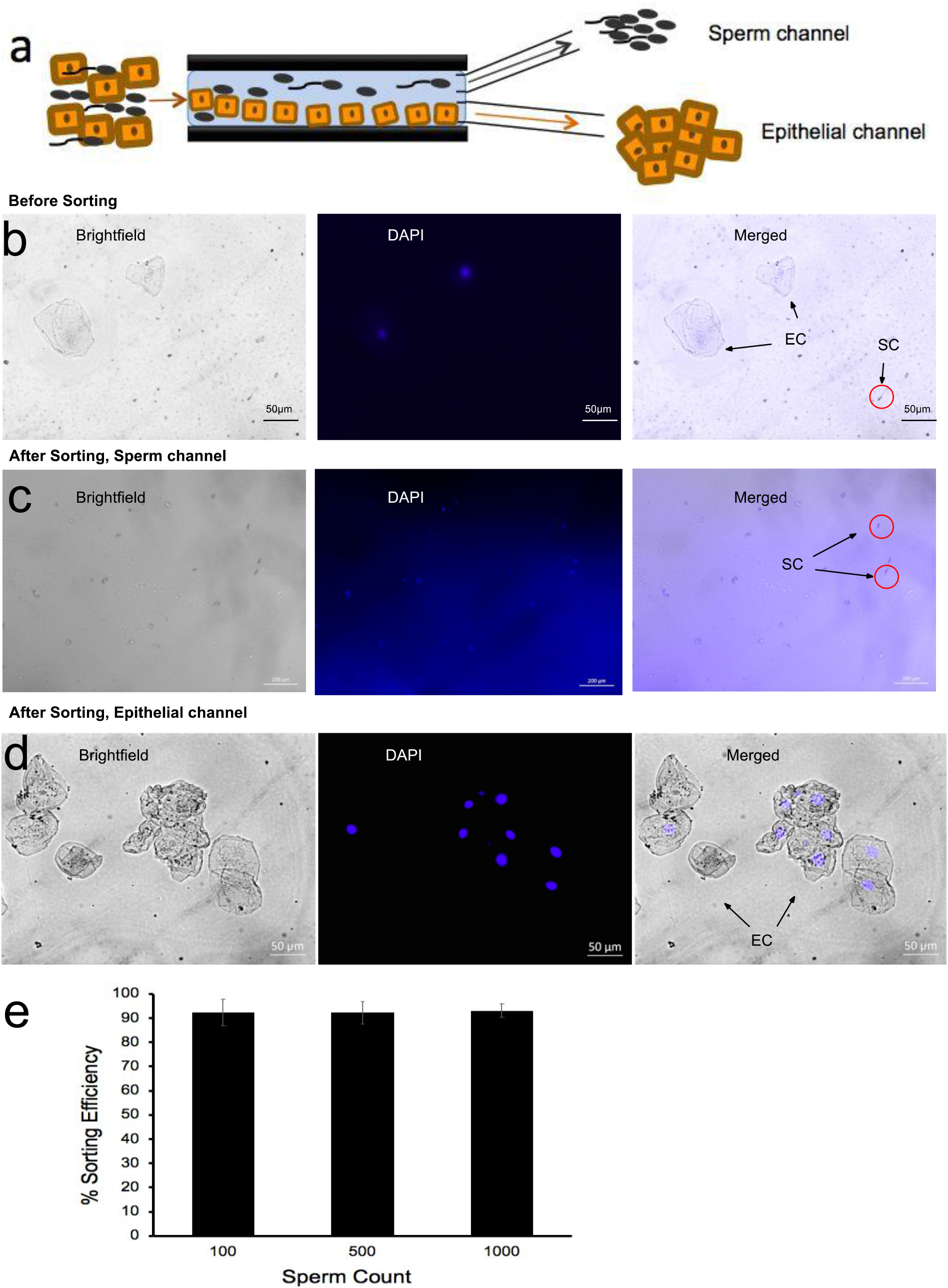
Sorting of sperm from epithelial cells. **a)** Schematic representation of the experimental workflow for spike-in experiments. 100, 500, 1,000 sperm cells/sample volume were spiked in 10,000 epithelial cells in 30 mM paramagnetic medium. The mixture was levitated and sperm were collected from the top channel (sperm channel), and epithelial cells were collected from the bottom channel (epithelial channel). **b)** Images of sperm and epithelial cell mixture before sorting are taken under bright field, DAPI, and merged. Sperm cells are shown with arrows as “SC” and epithelial cells are shown as “EC”. **c)** Images of the sperm sorted from the sperm channel under bright field, DAPI, and images are merged. Sperm cells can be detected as blue, since the spiked sperm were stained with DAPI. **d)** Images of the epithelial channel after sorting are taken under bright field, DAPI, and merged. Epithelial cells can be detected as blue, since the excess DAPI stained the epithelial cells during sorting. **e)** Sorting efficiency of 100, 500, 1000 sperm cells/sample volume in 10,000 epithelial cell sample is calculated. The efficiency is defined as the ratio of the number of sperm cells collected in the top channel and the number of sperm cells detected in the capillary.

### 2.4. Validating Magnetic Levitation Platform Performance with Forensic Mock Samples

To validate the method with samples that are most representative of a real case sample, we tested three mock forensic samples dating back to 2003 (Sample #1, 2), and 2015 (Sample #3). The collected samples were non-casework samples, including epithelial cells and sperm. They were low in volume representative of an actual forensic sample and were obtained from swab and gauze. Sperm cells tend to deteriorate quickly after ejaculation^[11]^. Tail of the sperm is more susceptible to damage and they break down first in the forensic samples^[12]^. Thus, the sperm found in forensic swab samples are mostly dead and non-motile, especially due to years of shelf-life after their collection from the crime scenes^[13]^. E-cells in forensic samples are mostly dead and their membrane integrity and permeability is compromised, leading the paramagnetic solution to rush inside the cell membrane^[9]^. This leads to a loss of levitation for these cells making the spatiotemporal separation for these cell types and the following sorting process easier for these samples. We observed that the sperm preserve their integrity and are not permeable to the paramagnetic solution and they overall levitate higher than the epithelial cells (Figure 5a). The forensic mock samples were imaged before sorting (Figure 5b), and after sorting in order to quantify the sorting efficiencies (Figure 5c, d). Despite the old age of the mock samples, cells kept their levitation profiles and their levitation heights were analyzed (Figure S2, Supporting Information). After injecting the sample into the levitation system, sperm and epithelial cell counts were recorded. The sample was then sorted. We calculated the sorting efficiency for sperm to be 90.4 ± 0.95% (n=3) (Figure 5e, Table S5, Supporting Information). Sorted cells kept their integrity although they were dead and fragile. In addition, we calculated the purity of the sperm sample collected from the top channel during sorting (Figure 5f, Table S6, Supporting Information). The purity of the sorted sperm from three different mock samples was 97.0%, 97.30% and 98.20%; respectively (n=3).

**Figure 5.**
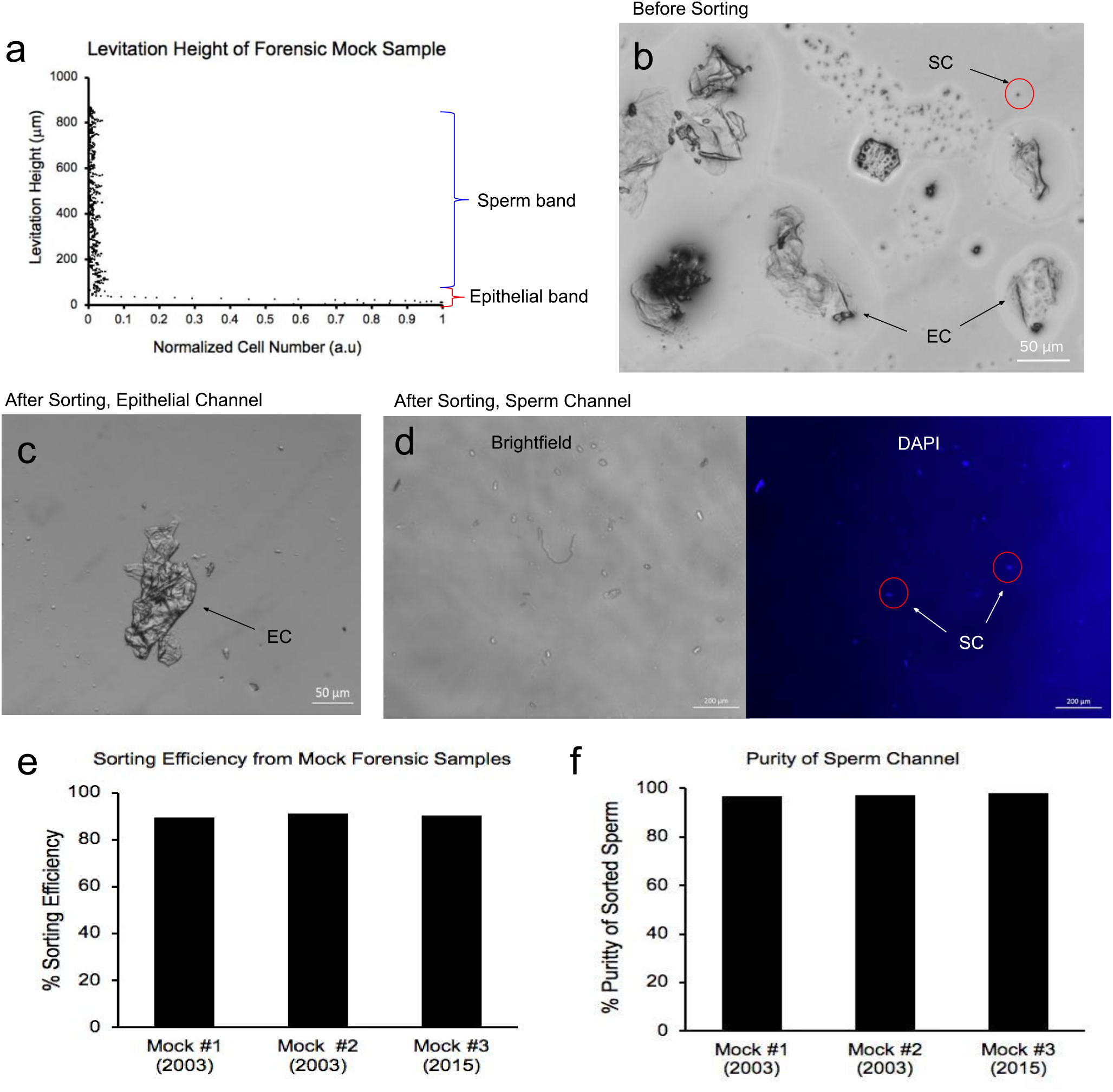
Sorting of forensic mock samples dating back to 2003. A mixture of epithelial and sperm cells eluted from mock swab samples were introduced into the magnetic levitation system. **a)** Average levitation height graphs of sperm and epithelial cells in the mock sample is plotted. Within minutes, epithelial cells sinked near to the bottom of the microcapillary and form a tight levitation band, while sperm cells form a dispersed levitation band along the microcapillary. **b)** A representative image of mock sample is taken before sorting. Epithelial cells are marked with arrows as “EC”, and sperm cells are marked as “SC” on the image. **c)** A representative image of the epithelial channel after levitational sorting. **d)** A representative image of the sperm channel taken after sorting under bright field and DAPI channel. **e)** Sorting efficiency of three mock samples. Sorting efficiency is defined as the ratio of the number of sperm cells detected in the sperm channel after sorting and the number of sperm cells detected in the capillary channel before sorting. Average sorting efficiency for three mock sample is 90.4% (n=3). **f)** Percent purity of the sperm channel is calculated for three mock samples after sorting. Percent purity is defined as the ratio of the number of epithelial cells detected in the sperm channel and the total number of cells detected in the sperm channel. The purity of the sorted sperm from three different mock samples is 97.0%, 97.30% and 98.20%; respectively (n=3).

## 3. Discussion

Here, we demonstrated that the sperm and epithelial cells can be detected under a magnetic field at different levitation heights. The long shelf-time of the forensic mock samples contribute to the epithelial cells to easily lose their membrane integrity and lose the ability to levitate easily. Sperm, on the other hand, is shown to keep this integrity over the years facilitating a simpler path to separation using mock samples. Thus, the broader sperm levitation band observed with mock samples can be attributed to the fact that sperm in these samples can become more of a heterogeneous population in terms of their levitation height. Even though this may be partly due to the jeopardized membrane permeability over years of shelf-life, sperm still levitate above the epithelial band at concentrations optimized for levitation. This is further signified with the dead epithelial cells that can levitate above the channel bottom at much higher paramagnetic concentrations, although they are permeable to the paramagnetic solution. Even though this may be partly due to the jeopardized membrane permeability over years of shelf-life, sperm still levitate above the epithelial band at the paramagnetic medium concentrations optimized for levitation. This is further signified with the dead epithelial cells that can levitate above the channel bottom at much higher paramagnetic concentrations, although they are permeable to the paramagnetic solution. This observation indicates that the membrane integrity is not necessary for a levitation event to enable differential sorting of dead cells, which cannot be achieved by any other tool otherwise. Further, epithelial cells are also recovered from the bottom channel as they might carry forensic value in specific cases. Further, epithelial cells are also recovered from the bottom channel as they might carry forensic value in specific cases.

Overall, we presented that the levitation of cells as a method provides a rapid, label-free, inexpensive and high-yield alternative to current analysis methods. The outcome of our novel cell separation methodology can open up new avenues for downstream analyses in forensics impacting the analysis of backlogged sexual assault cases and as a promising technology for the magnetic levitation-based separation of heterogeneous biological fluids.

## 4. Experimental Section

### Fabrication of Magnetic Levitation Platform and Flow Setup

A magnetic levitation chip consists of polymethyl methacrylate (PMMA), permanent neodymium magnets and aluminum-coated mirrors. A microchannel with 1-mm x 1-mm cross-section was placed between two permanent neodymium magnets (NdFeB) with same poles facing each other. For high-resolution spatiotemporal and real-time imaging purposes, two mirrors were placed at each side of the microchannel^[9]^. The levitation chip can be placed under either a 5X or 20X objective on the microscope and levitating cells (i.e., sperm, epithelial cells) can be imaged. The optical path to observe the cells under the magnetic field is shown in Supplementary Figure S3. The flow-based magnetic levitation system was driven by syringe pumps that withdrew the sample into the levitation chip. The outlet flow from the magnetic levitation chip was controlled by two different syringe pumps: The top outlet was connected to a pump that withdraws the solution containing the sperm; while the bottom outlet was connected to a pump that withdraws the solution containing the epithelial cells. In all sorting experiments, a 4:3 flow ratio was maintained between the top and bottom outlet flow streams.

### Preparation of Sperm Samples

Semen samples were purchased from California CryoBank under an Institutional Review Board (Stanford University IRB Number: 6208, and Protocol ID: 30538). The semen sample was centrifuged at 5,000 rpm for 5 minutes. After centrifugation, the liquid was discarded, and the resulting sperm pellet was re-suspended in PBS. 10 μL of DAPI stained was put in 2 mL of sample. After a 15-minute incubation, the sample was centrifuged at 5,000 rpm for 5 minutes. The excess stain was disposed. The sperm sample was then diluted with 1:5 dilution ratio, and counted for cells/mL with the use of hemocytometer.

### Preparation of Epithelial Samples

Buccal epithelial cells were collected from female individuals. Briefly, a cotton swab was utilized to capture the epithelial cells. The swab was then suspended in PBS for 15 minutes. The sample was centrifuged at 5,000 rpm for 5 minutes. The liquid at the top layer was disposed, and the cell pellet was re-suspended in PBS. The sample was then diluted with a 1:10 dilution ratio. Cells/mL count was obtained with the use of a hemocytometer.

### Levitation of Sperm

Known amounts of sperm (100 sperm, 500 sperm, 1,000 sperm) were suspended in different paramagnetic concentrations (i.e., 30 mM, 100 mM). The volume of the mixture was set to 30 μL with the addition of PBS. The mixture was then pipetted in the microchannel. The samples were loaded in the magnetic levitation chip and levitated for 15 minutes. Levitation heights of sperm were imaged and quantified with the in-house developed image analysis software. The detection efficiency was calculated by counting the number of DAPI-positive sperm seen in the capillary.

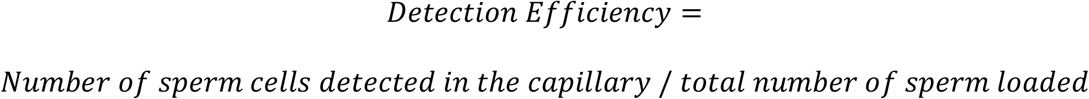

### Levitation of Epithelial Cells

Known number of epithelial cells (i.e., 10,000 cells, 50,000 cells) was mixed with known concentration of paramagnetic medium (i.e., 30 mM, 100 mM). The volume of the mixture was set to 30 μL with the addition of PBS. The mixture was then loaded in the magnetic levitation chip and levitated for 5 minutes. Levitation heights of epithelial cells were imaged and quantified with the in-house developed image analysis software.

### Spiking of Sperm in Epithelial Cells for Separation Efficiency Characterization

The separation efficiency of the static magnetic levitation chip was tested using epithelial cells spiked with a known number of sperm (100 sperm, 500 sperm, and 1,000 sperm/channel). 30 μl of epithelial cell + sperm mixture was pipetted into the microchannel. Then, the samples were loaded into the magnetic levitation chip and were levitated for 10 minutes. Levitation heights of separated sperm and epithelial cell populations were imaged and quantified with the in-house developed image analysis software.

Separation efficiency is defined as the ratio of sperm levitated above the epithelial cells to the total number of spiked sperm. Experiments were performed in triplicates and repeated at least three times to validate repeatability and reproducibility. Data are represented as the mean ± standard error of the mean (SEM).

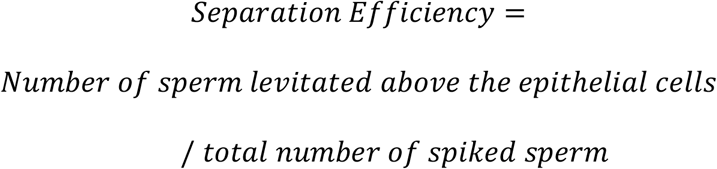

### Sorting Efficiency of Sperm Cells from Epithelial Cells

After running the sample, a washing step with PBS was utilized in order to minimize the loss of sperm due to their adherence to the inlet and outlet tubes. Then, the collected samples from the top channel and bottom channel in well-plates were centrifuged at 2,000 rpm for 5 minutes. We imaged each well to count the sperm and epithelial cells. Sorting efficiency was calculated for each sample according to the equation below.

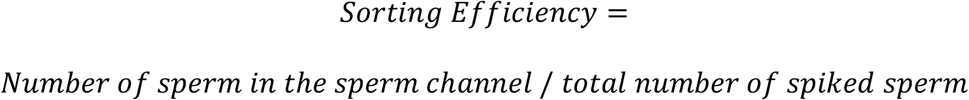

### Forensic Mock Samples

Mock samples were prepared in collaboration with Broward Sheriff’s Office Crime Laboratory. Cotton swabs containing simulated forensic samples were placed in 500 μL of 1X PBS and mixed at 4°C in a Thermomixer (Eppendorf, Germany) set at 1,000 rpm for approximately an hour. Then, the cotton cuttings were removed and centrifuged in spin baskets at 16,100 rcf/13,200 rpm for 5 minutes to pellet the solids in the solution. After using pulse vortexing to re-suspend the pellet, 5 μL of each sample was placed on a glass slide and imaged for confirmation before levitational sorting experiments. The forensic mock sample experiments were processed with a 10 μL sample volume at the earlier optimized 30 mM paramagnetic concentration. After sorting, sorting efficiency and percent purity of the sperm channel was calculated according to the equation below.

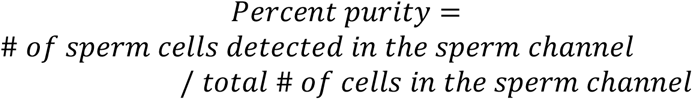

### Image Analysis of Levitated Samples

Image analysis and characterization of levitation profiles was performed using a customized in-house developed code, employing the Image Processing toolbox in MATLAB. The image datasets were acquired using the ZenPro2 software (Zeiss) and were imported into the code in “.tiff” format.

## Acknowledgements

N.G.D acknowledges support from the Career Award at the Scientific Interface (CASI) from the Burroughs Wellcome Foundation (BWF). N.G.D. acknowledges support from the McCormick and Gabilan Faculty Award from Stanford University.

