## Supplementary Information for "Levitational Cell Cytometry for Forensics"

**Underlying mechanisms for levitation of cells in a microcapillary:**
The magnetic susceptibility difference between a cell and its surrounding paramagnetic medium causes it to move away from a higher to a lower magnetic field strength site until gravitational, buoyancy and magnetic forces acting on the cells reach an equilibrium. Cells are levitated at a final position, where the magnetic force (Fmag) equals the buoyancy force (Fb). As the cells flow, the platform monitors the equilibrium heights of each individual cell in real-time, magnetically focuses and separates sperm and epithelial cells at different levitation bands, based on their unique magnetic signatures. We have shown that the epithelial cells get suspended in a higher levitation height when 100 mM Gd. concentration is used. Thus, we can conclude that the levitation height of epithelial cells at the bottom of the channel under 30 mM Gd. concentration isn't solely due to the gravity but is rather caused by magnetic forces in the levitation device.

We also provide here a more description of the physical support explaining the principles of magnetic levitation of cells. During levitation, magnetic force ($\mathbf{F_{mag}}$), buoyancy force ($\mathbf{F_b}$) and drag forces ($\mathbf{F_d}$) are induced on the cells can be calculated:

$$F_{mag} + F_b + F_d = 0 \qquad \text{(Eq.1)}$$

Cells are levitated in the channel with $\mathbf{F_{mag}}$ (42):

$$F_{mag} = (m.\nabla)B \qquad \text{(Eq.2)}$$

where **B** is the magnetic induction, $\nabla$ is the del operator and **m** is the magnetic moment, which is calculated as:

$$m = \frac{V\Delta\chi}{\mu_0}B \qquad \text{(Eq.3)}$$

with $V$ the volume of the cell, $\mu_0$ the permeability of the free space ($1.2566\times10^{-6}$ kg m $A^{-2}$ $s^{-2}$) and $\Delta\chi$ the magnetic susceptibility difference between the cell and paramagnetic medium. **B** induced in the channel by opposing magnets in the levitation setup.

During levitation, cell gained velocity **v** and $\mathbf{F_d}$ is exerted on the cell. $\mathbf{F_d}$ is calculated for spherical object as follows (42):

$$F_d = 6\pi R\eta f_D v \qquad \text{(Eq. 4)}$$

,where $R$ is the radius of the cell, $\eta$ is the dynamic viscosity of the paramagnetic medium and $f_D$ is the drag coefficient, which is equal to 1 when the cell is far away from the channel wall.

$\mathbf{F_b}$ is calculated as (44):

$$F_b = V\Delta\rho g \qquad \text{(Eq. 5)}$$

,where **g** is the gravitational acceleration (9.8 $ms^{-2}$), in z-direction, and $\rho$ is the difference between the volumetric densities of the cell and the paramagnetic medium.

In the levitation setup, cells are focused on $x=0$ plane where $B_x$=0 with magnetic forces. On the other hand, the cell levitates in a certain height in **z** direction along $x=0$ plane until magnetic and buoyancy forces come into a balance. During levitation, the equilibrium height of a cell is calculated based on magnetic induction values $B$ (24-26):

$$F_{mag} + F_b = 0 \qquad \text{(Eq. 6)}$$

$$\frac{\Delta\chi}{\mu_0}\left(B_x\frac{\partial B_z}{\partial x} + B_y\frac{\partial B_z}{\partial y} + B_z\frac{\partial B_z}{\partial z}\right) - \Delta\rho g = 0 \qquad \text{(Eq. 7)}$$

The gravitational acceleration is represented by $g$, $\mu_0$ defines the permeability of the free space, $\nabla\rho$ is the volumetric density difference between cell and paramagnetic medium (*i.e.*, $\rho_c$-$\rho_m$) and $x$, $y$, and $z$ are the coordinates of a cell. The magnetic susceptibilities of cells (27) are negligible compared to the magnetic susceptibility of the paramagnetic medium used for levitation (28). Thus, cells are equilibrated at a unique levitation height mainly based on their density, independent of their volume. For instance, cells with the same density as the paramagnetic medium are equilibrated in the middle of the channel and cells with different densities than the medium are equilibrated above (if $\rho_c<\rho_m$) or below the middle of the channel (if $\rho_c>\rho_m$). In addition, cells are focused along the $x$-axis towards the middle of the channel, where the magnetic induction strength is lowest, as we have shown in our earlier work.

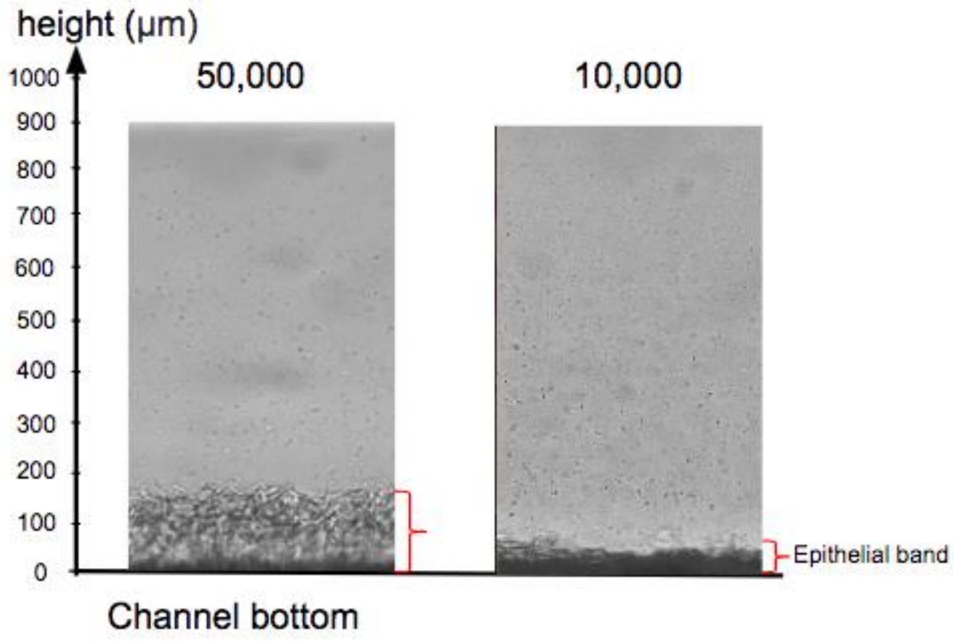


**Supplementary Figure S1. Levitation profiles of 10,000 and 50,000 epithelial cells/sample volume.** Levitation profiles for 50,000 and 10,000 cells/sample volume are imaged in 30mM paramagnetic medium concentration. It is shown that 10,000 epithelial cells have a thinner levitation band than that of 50,000 epithelial cells.

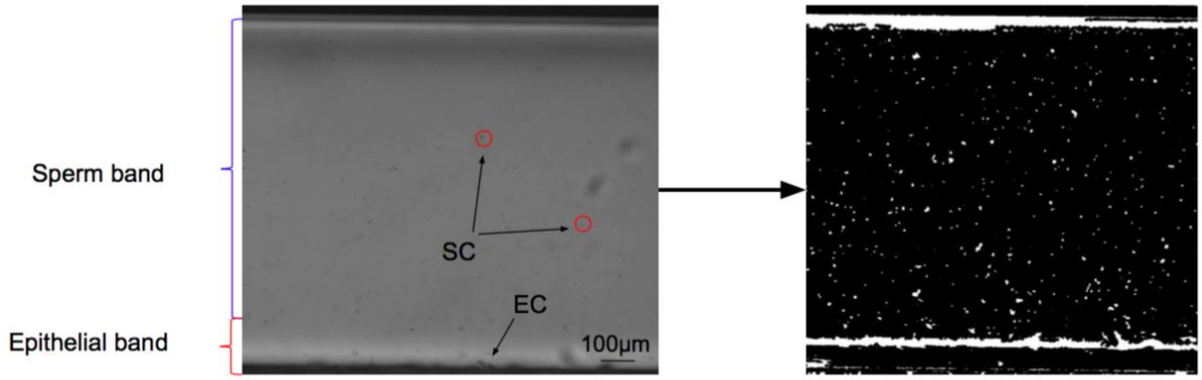


**Supplementary Figure S2. Levitation and image analysis of mock samples.** Mock samples were levitated in 30mM paramagnetic medium. Sperm cells are shown as “SC” and epithelial cells are shown as “EC” in the first image. Then, the levitation profiles were analyzed using MATLAB single cell detection code, in order to plot the average levitation height graphs of the two cell types.

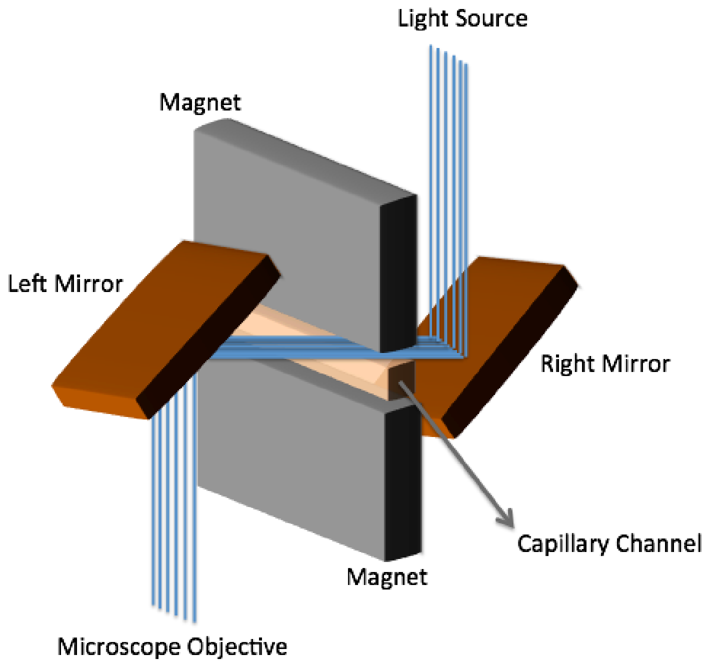


**Supplementary Figure S3. Optical Setup.** Two aluminium-coated mirrors are placed at each open side of the glass microcapillary tilted at 45 degrees to make the levitation platform compatible with conventional microscopy systems for high resolution spatiotemporal monitoring of cells during levitation.

**Supplementary Table S1. Comparison of magnetic levitation platform with other methods for sorting and extraction of sperm.**

| Methods | Needed sperm concentration | Labels | Sperm sorting/detection accuracy | Assay time |
|---|---|---|---|---|
| Magnetic levitation method | $>10^2$ sperm per capillary channel | Label-free | >90% with samples dating back to 2003 | 20 min. |
| Magnetic bead-based separation ([5]) | $>10^4$/mL | Immunomagnetic beads | 16.67% for swabs preserved for 10 days | N/A |
| Acoustic differential extraction ([3]) | $>2 \times 10^4$/μL | Label-free | N/A | 14 min. |
| On-Chip Method ([6]) | $>10^2$ sperm per channel | SLeX | 70–92% | 80 min. |

**Supplementary Table S2. Sperm detection efficiency.** Sperm detected in each trial of 100, 500, and 1,000 sperm/sample volume is recorded to calculate the percent yield.

| Number of cells loaded | 1st Trial count | 2nd Trial count | 3rd Trial count | Average count | Percent Yield |
|---|---|---|---|---|---|
| 100 cells | 100 | 80 | 100 | 93.3 | 93.3% |
| 500 cells | 495 | 480 | 494 | 489.6 | 97.9% |
| 1,000 cells | 992 | 920 | 874 | 928.0 | 92.8% |

**Supplementary Table S3. Separation efficiency of sperm from epithelial cells.** Separation efficiency of each trial is recorded to get an average separation efficiency for different sperm concentrations.

| Cell number | Separation efficiency Trial 1 | Separation efficiency Trial 2 | Separation efficiency Trial 3 | Average |
|---|---|---|---|---|
| 100 sperm & 10,000 epithelial | 100.0% | 90.0% | 91.8% | 93.9% |
| 500 sperm & 10,000 epithelial | 92.8% | 97.5% | 97.8% | 96.1% |
| 1,000 sperm & 10,000 epithelial | 90.0% | 95.7% | 97.6% | 94.4% |

**Supplementary Table S4. Sorting efficiency of spiked sperm from epithelial cells.** Sorting efficiency of each trial is recorded to get an average sorting efficiency for different sperm concentrations.

| Cell number | Sorting efficiency Trial 1 | Sorting efficiency Trial 2 | Sorting efficiency Trial 2 | Average sorting efficiency |
|---|---|---|---|---|
| 100 sperm & 10,000 epithelial | 100.0% | 82.0% | 90.0% | 90.7% |
| 500 sperm & 10,000 epithelial | 100.0% | 92.6% | 84.0% | 92.2% |
| 1,000 sperm & 10,000 epithelial | 89.5% | 98.5% | 91.2% | 93.6% |

**Supplementary Table S5. Sorting efficiency of forensics mock samples.** Sorting efficiency of mock samples is recorded for each trial.

| Sorting efficiency Sample 1 | Sorting efficiency Sample 2 | Sorting efficiency Sample 2 | Average sorting efficiency |
|---|---|---|---|
| 89.4% | 91.3% | 90.5% | 90.4% |

**Supplementary Table S6. Percent purity of the sperm channel of mock samples.** Percent purity of sperm channel is calculated from the ratio of the number of sperm and the number of total cells collected in the sperm channel.

| Sample number | E. cells in sperm channel | Sperm in sperm channel | Percent purity of sperm channel |
|---|---|---|---|
| #1 | 5 | 160 | 97.0% |
| #2 | 4 | 146 | 97.3% |
| #3 | 3 | 165 | 98.2% |

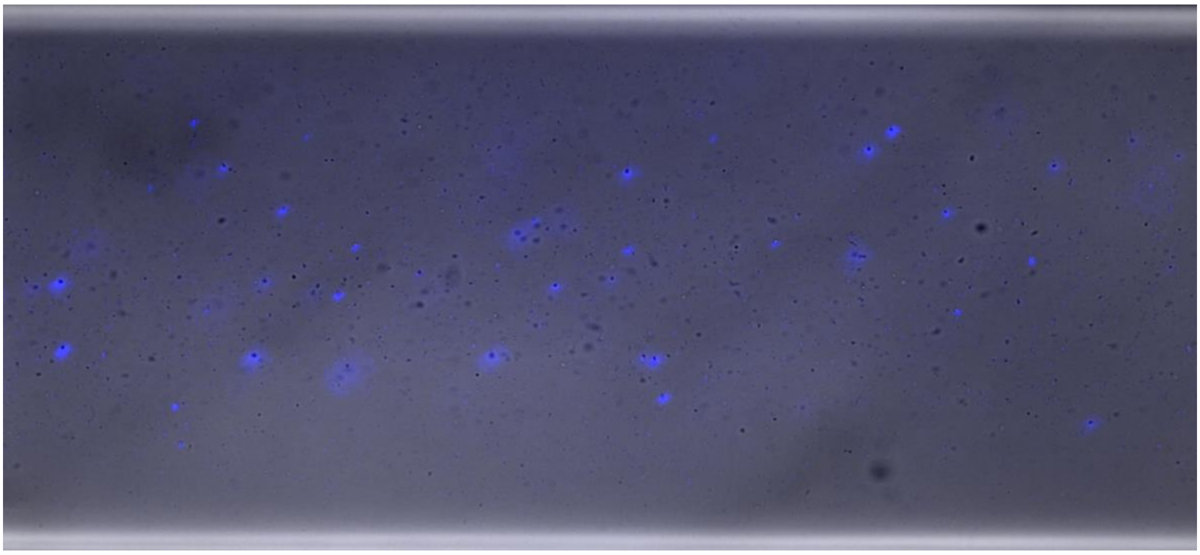

**Supplementary Video S1.** Static levitation of 1,000 sperm stained with DAPI in 30 mM paramagnetic medium.

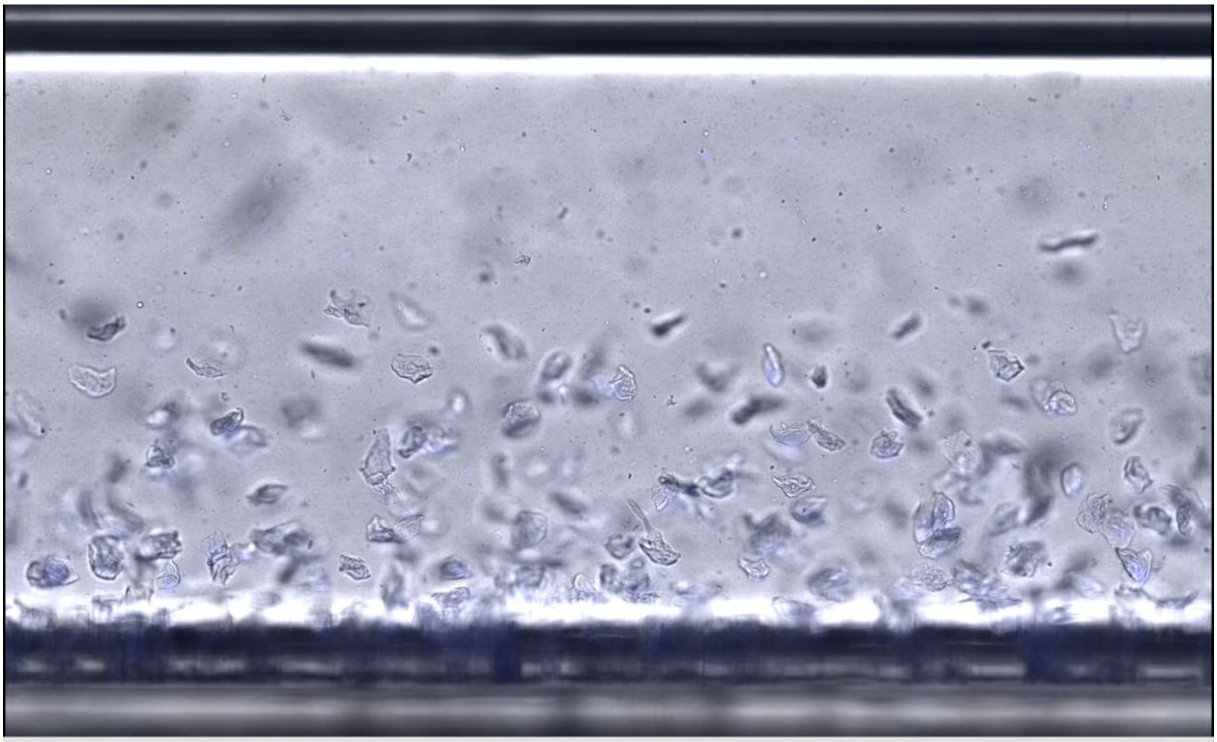

**Supplementary Video S2.** Static levitation of 1,000 sperm and 10,000 epithelial cells in 30 mM paramagnetic medium.

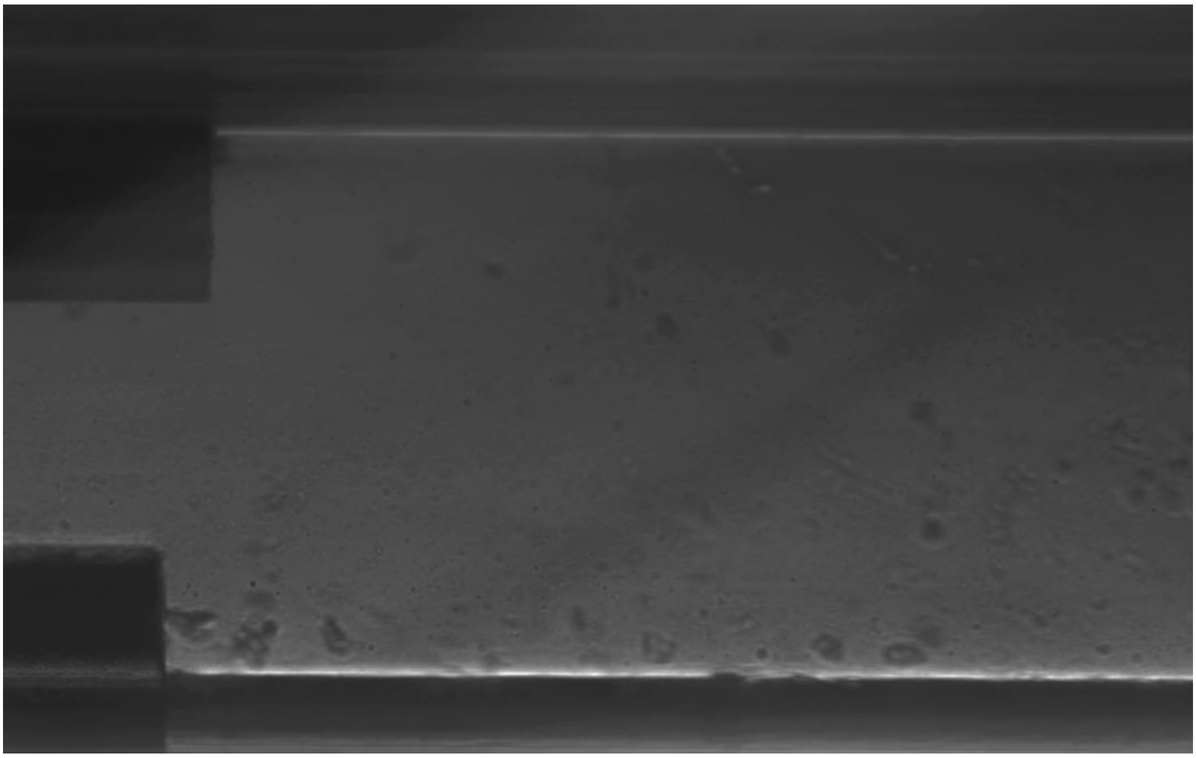

**Supplementary Video S3.** Sorting of 1,000 sperm and 10,000 epithelial cells in 30 mM paramagnetic medium. Epithelial cells are sorted to the bottom collection channel, whereas the sperm are sorted to the top collection channel.

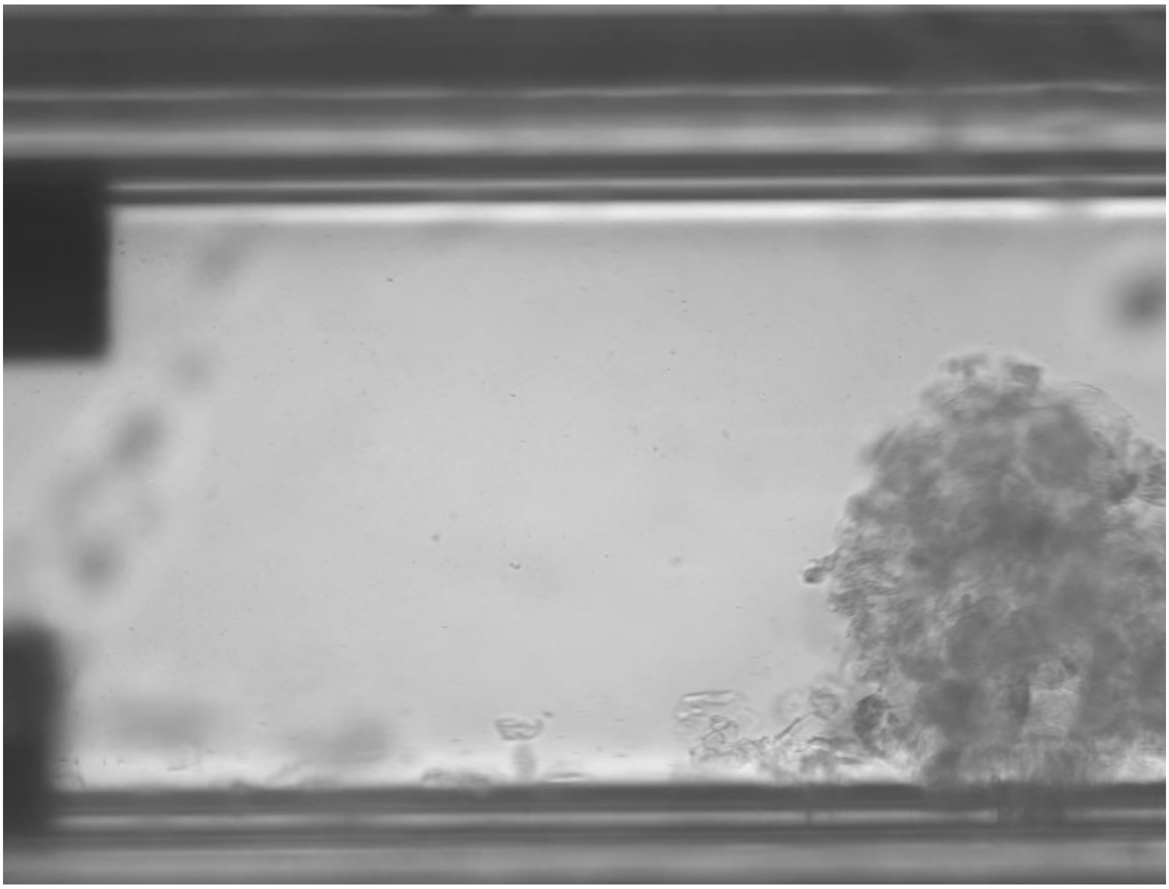

**Supplementary Video S4.** The end of sorting of 1,000 sperm and 10,000 epithelial cells in 30 mM paramagnetic medium.
